# Co-expression analysis is biased by a mean-correlation relationship

**DOI:** 10.1101/2020.02.13.944777

**Authors:** Yi Wang, Stephanie C. Hicks, Kasper D. Hansen

## Abstract

Estimates of correlation between pairs of genes in co-expression analysis are commonly used to construct networks among genes using gene expression data. Here, we show that the distribution of such correlations depend on the expression level of the involved genes, which we refer to this as a *mean-correlation relationship* in RNA-seq data, both bulk and single-cell. This dependence introduces a bias in co-expression analysis whereby highly expressed genes are more likely to be highly correlated. Such a relationship is not observed in protein-protein interaction data, suggesting that it is not reflecting biology. Ignoring this bias can lead to missing potentially biologically relevant pairs of genes that are lowly expressed, such as transcription factors. To address this problem, we introduce spatial quantile normalization (SpQN), a method for normalizing local distributions in a correlation matrix. We show that spatial quantile normalization removes the mean-correlation relationship and corrects the expression bias in network reconstruction.

## Introduction

Gene co-expression analysis is the study of correlation patterns in gene expression data, usually with the goal of constructing gene networks. Amongst popular methods for gene co-expression analysis is WGCNA (Langfelder, Horvath, 2008) which works on the correlation matrix and the graphical LASSO (Friedman et al., 2008) which works on the precision matrix, the inverse of the co-variance matrix. These methods have been successfully used many times to gain biological insight (Ghazalpour et al., 2006; Oldham et al., 2008; Willsey et al., 2013; Saha et al., 2017; Boukas et al., 2019).

While co-expression analysis is widely used, there has been less work on various sources of confounding and bias, especially in contrast to the rich literature on such issues in the related field of differential gene expression analysis. Recent work in co-expression analysis is addressing this shortcoming, including work on removing unwanted variation (Freytag et al., 2015; Parsana et al., 2019) and the effect of cell type composition (Zhang et al., 2019).

Recently, it was shown that differential expression confounds differential co-expression analysis (Farahbod, Pavlidis, 2019). Defining differential co-expression as changes in correlation patterns between conditions, the authors show that most correlation changes are associated with changes in gene expression levels in the same genes between condition, but provide no method to address this confounding. This work highlights the importance of considering changes in expression level when interpreting changes in correlation patterns. Note however, that co-expression is commonly performed within a single condition.

## Results

### The distribution of gene-gene correlations depend on gene expression level

We have previously observed that highly expressed genes tend to be more highly co-expressed (Boukas et al., 2019). Related, differential expression has been shown to confound differential co-expression (Farahbod, Pavlidis, 2019). Here, we use bulk RNA-seq data from 9 tissues from the GTEx project (GTEx Consortium, 2017) to further explore the relationship between gene expression and gene-gene co-expression.

Gene counts were converted into log-RPKMs (Methods). Starting with 19,836 protein-coding genes and 7,036 lncRNAs, in each tissue we kept genes with a median expression above zero (on the log-RPKM scale, Methods), leaving us with 10,735-12,889 expressed genes per tissue (of which 95%-98% were protein-coding). Removing principal components from the correlation matrix has been shown to remove unwanted variation in co-expression analysis (Parsana et al., 2019). We computed the gene-gene correlation matrix of the log-RPKM values and removed unwanted variation by removing the top 4 principal components (PCs). The number 4 was chosen based on our previous analysis (Boukas et al., 2019), where we used positive and negative control genes to assess this. An alternative approach suggested by Parsana et al. (2019) is to remove the number of PCs according to the estimated number of surrogate variables using SVA (Leek, Storey, 2007; Leek, Storey, 2008; Leek, Johnson, et al., 2012) with the number of PCs ranging from 10 to 30 in these same tissues – the impact of choosing a different number of PCs will be examined below.

For each tissue, we sorted genes according to their expression level (average log-RPKM across replicates of that tissue) and grouped them into 10 bins of equal size. We number these gene expression bins from 1 to 10, with 1 being lowest expression. Then, we divided the gene-gene correlation matrix into a 10×10 grid of 100 non-overlapping submatrices numbered as (*i, j*) using the expression level for the *i*^*th*^ and *j*^*th*^ bin (Figure 1). The use of a 10×10 grid is somewhat arbitrary, but it ensures a substantial number of correlations inside each submatrix.

**Figure 1.**
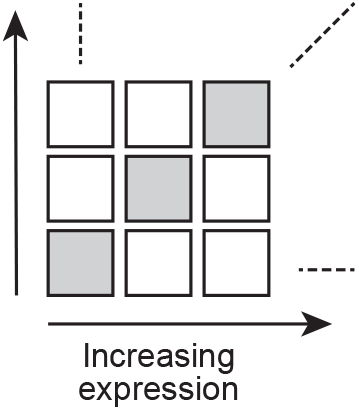
Partitioning the correlation matrix. Genes are ordered according to expression level, and the correlation matrix is partitioned into 10×10 non-overlapping submatrices. The diagnonal submatrices are indicated by grey.

As an example, we focus on adipose subcutaneous. For this tissue, if we focus on the distribution of correlations within the 10 diagonal submatrices, we observe that these distributions are all centered around 0, but their variance increase with expression level (Figure 2a). A robust estimate of the spread of the distribution is given by the interquartile-range (IQR), the difference between the 25% and 75%-quantiles. We can depict the IQR across the binned correlation matrix, forming what we term a 2D boxplot (Figure 2b). This reveals the *mean-correlation* relationship: namely, the IQR of each submatrix is dependent on the expression level of the two associated bins, and that the IQR is approximately dependent on the minimum of the two expression levels (Figure 2c).

**Figure 2.**
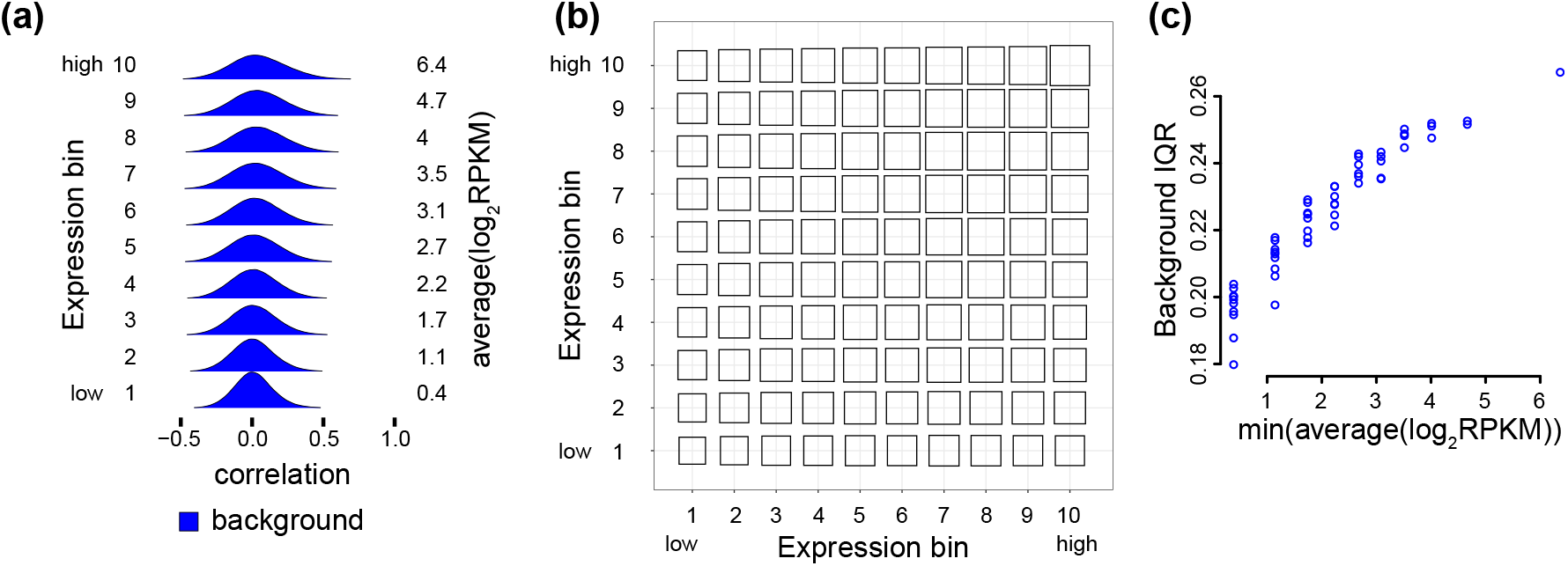
The mean-correlation relationship between gene expression level and the distribution of observed gene-gene correlations. The distribution of Pearson correlations of gene pairs using 350 RNA-seq samples from adipose subcutaneous tissue, with 4 PCs removed. **(a)** Densities of the Pearson correlation between gene pairs stratified by overall expression (10 bins ranging from low to high expression). Average expression level for each expression bin is given by the values to the right of the densities. **(b)** A 2D boxplot where each box represents the IQR of the Pearson correlations between all genes (termed background IQR) in a submatrix of the correlation matrix corresponding to two bins of expression. **(c)** The relationship between IQRs of the Pearson correlations between all genes in a submatrix, and the minimum of the two expression bins associated with the submatrix.

In Freytag et al. (2015), it is suggested that genes selected at random should be uncorrelated, and the authors verify this to be true empirically, in multiple datasets. Another way of stating this assumption is that the true gene-gene correlation matrix of random genes should be sparse. Therefore, under this model we expect that the observed distribution of gene-gene correlations to be made up of mostly true correlations of zero coupled with some “background” (or random) noise centered around 0. This is assumption is supported by the observed distributions of pairwise correlations depicted in Figure 2, which are largely symmetric around 0 – one or two expression bins have a slight location shift away from 0. However, in Figure 2 we also see that the background distribution depends on gene expression level.

### The mean-correlation relationship biases co-expression analysis

In co-expression analysis the goal is often to identify biologically meaningful gene pairs with what is assumed to be high correlations (signal) from ones with random gene pairings with usually low correlation (noise). Therefore, a common first step is to separate these highly correlated gene pairs, or clusters (sometimes called modules), from the lowly correlated gene pairs. There are multiple approaches, including thresholding the correlation matrix, using weighted gene correlation analysis (WGCNA) (Langfelder, Horvath, 2008) or using the graphical LASSO, which operates on the precision matrix; the inverse of the covariance matrix (Friedman et al., 2008).

To visualize the (possible) signal component of the correlation matrix, we overlay the full (background) distribution with the distribution of the top 0.1% of the correlations (signal) within each expression bin. In addition to the mean-correlation relationship in the background distribution, we see that the signal component of the correlation distributions is also dependent on the expression level (Figure 3a). We observe that – conditional on the expression level – the signal and background distributions are well separated, but the point of separation between the two distributions depend on the expression level. As a consequence, thresholding the correlation matrix will result in an over-representation of highly expressed genes (Figure 3b), which explains our previous observation that highly expressed genes tend to be more highly co-expressed (Boukas et al., 2019). Here, we have used the average expression level of the two genes involved in a correlation, as a measure of the expression level of the pair.

**Figure 3.**
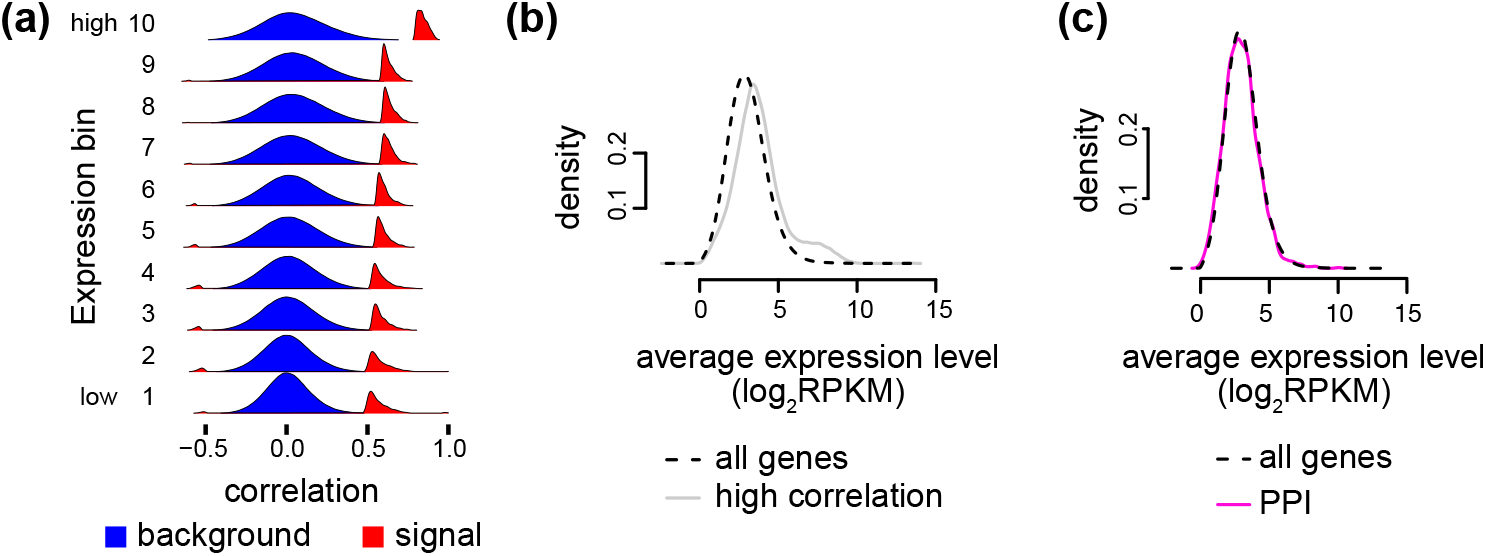
The mean-correlation relationship leads to expression bias. Same data as Figure 2. **(a)** Like Figure 2a but supplemented with the densities (scaled differently from the background densities) of the top 0.1% of the correlations in each bin, representing possible signal. **(b)** The expression level of pairs of genes for either all expressed genes (black) or all gene pairs in the top 0.1% of the correlations (gray). **(c)** The expression level of pairs of genes for either all genes (black) or all gene pairs with a known protein-protein interaction (PPI) (pink).

Furthermore, we do not observe this bias towards high expression when examining the expression level of gene pairs which are involved in known protein-protein interactions (PPIs) (Figure 3c). This strongly suggests that the observed bias is not biological and is unwanted.

### Model-based motivation for the mean-correlation relationship in RNA-seq data

A standard model for RNA-seq data is that we observe counts *ϒ* which reflect the true, but unknown, expression level *Z* > 0 and where *ϒ* | *Z* ~ Poisson(*Z*). Here, *ϒ* represents the expression level filtered by Poisson noise which we believe arise from the counting aspect of sequencing. This model has been experimentally verified (Marioni et al., 2008; Bullard et al., 2010). It is instructive to reflect on how this model can be extended to correlations.

Let *ϒ*_1_, *ϒ*_2_ be the data from two observed genes and *Z*_1_, *Z*_2_ be the unobserved “true” expression variables. We assume that E(*ϒ* | *Z*) = *Z*. Under mild assumptions, we can show (Methods)

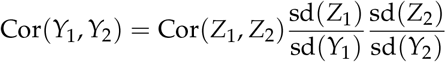

This expresses that the observed correlation is equal to a scaled version of the “true” correlation, with a gene-dependent adjustment factor. The adjustment factor is essentially driven by how much extra variation *ϒ* introduces on top of *Z*. Considering the adjustment factor, we can show (Methods)

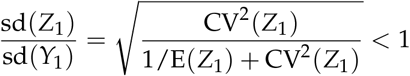

with CV^2^(*Z*_1_) being the squared coefficient of variation. This shows that the observed correlations are strictly smaller than the true correlations, with an adjustment factor close to 1 when the expression level of both genes is high.

This model explains why the width of the background distributions decrease with decreasing expression level (the adjustment factors decreases) and suggests that the “true” width of the background distribution is observable for highly expressed genes. Furthermore, it suggests that the background distributions in different sub-matrices are roughly related through a scaling transformation.

To explore whether the background distributions from different submatrices are related by scale transformations, we use quantile-quantile plots (Q-Q plots). If two distributions are related by a scale transformation, the Q-Q plot will be a straight line, with the slope of the line giving the scale parameter. Figure 4 suggests that a subset of the submatrices exhibit scaling differences, but that this is not true across all submatrices. The submatrices with non-linear differences are all at the “boundary” of the correlation matrix, with either very lowly expressed or very highly expressed genes (with a greater proportion of lowly expressed genes exhibiting this behavior). This result further motivated us to develop a method beyond scale transformations to correct for the mean-correlation bias.

**Figure 4.**
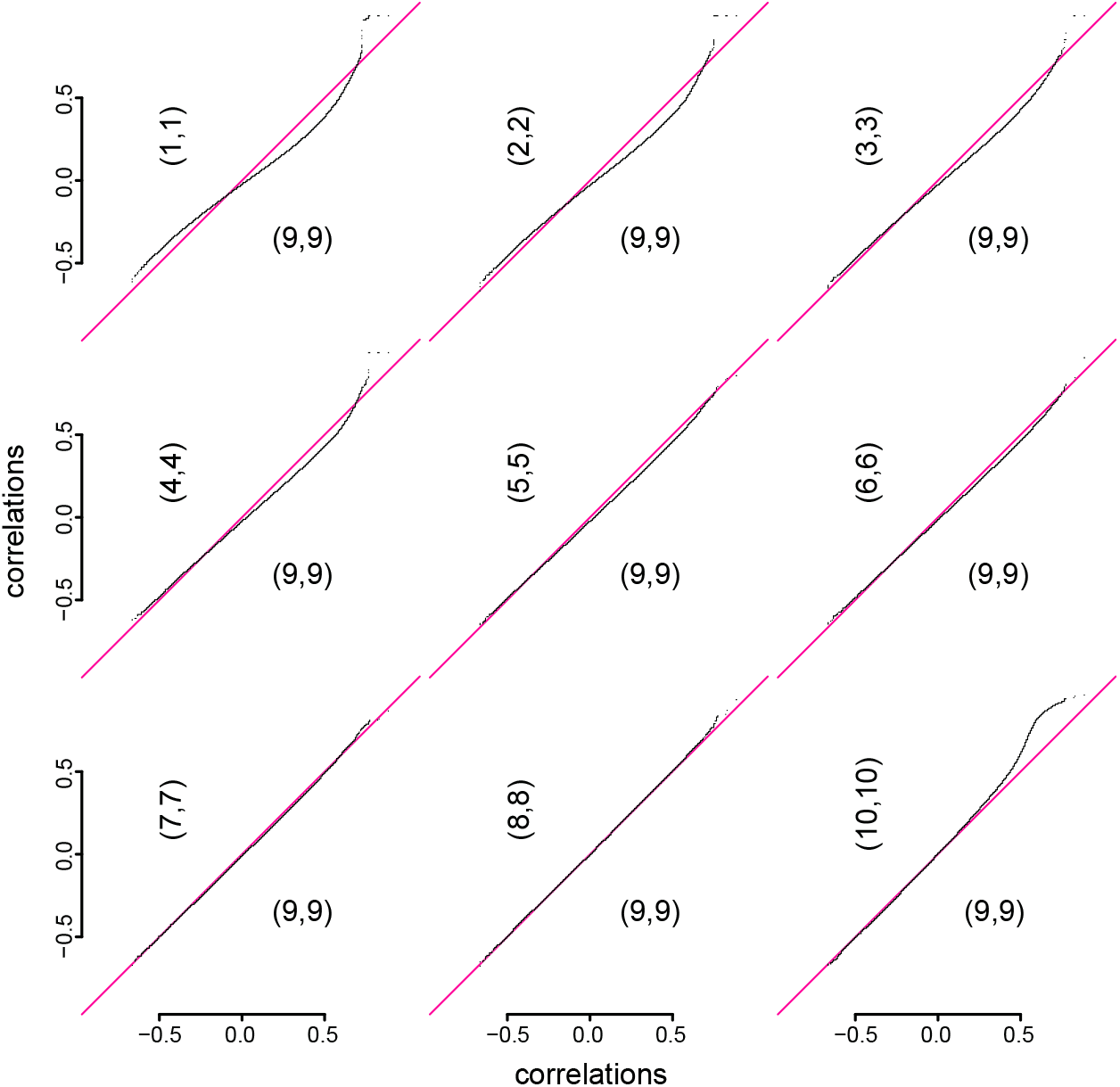
Distribution comparison for different submatrices of the observed correlation matrix (after removing the top 4 PCs) Same data as Figure 2. Quantile-quantile plots comparing the distribution of Pearson correlations in various (*i*, *i*) submatrices to the (9, 9) submatrix.

### Spatial quantile normalization

To correct for the mean-correlation bias, we developed a spatial quantile normalization method, referred to as SpQN. Here “spatial” refers to the spatial ordering of expression along the two dimensions of the correlation matrix. The objective is to achieve the same “local” distribution of correlations across the matrix. In other words, different submatrices should exhibit similar distributions. However, unlike quantile normalization (Amaratunga, Cabrera, 2001; Workman et al., 2002; Bolstad et al., 2003), our method does not mathematically guarantee that different submatrices end up with the same empirical distribution, although our experiments suggests that this is approximately true.

To explain our approach, recall that standard quantile normalization works by transforming observed data *X* using

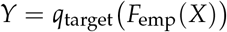

where *F*_emp_ is the empirical distribution function for *X* and *q*_target_ is the quantile function for a suitably chosen target distribution. In its original formulation of quantile normalization, *q*_target_ was chosen empirically as the average quantile distribution across samples.

Consider a submatrix *X*_*i,j*_ of the gene-gene correlation matrix (from the *i*^*th*^ and *j*^*th*^ ordered expression bins). Instead of using the empirical distribution (*F*_emp_(*X*_*i,j*_)) of *X*_*i,j*_ to form a distribution function – as is done in standard quantile normalization – we use a larger submatrix *ϒ*_*i,j*_ enclosing *X*_*i,j*_ as the basis of the empirical distribution function (Figure 5). This implies that when two sub-matrices, *X*_*i,j*_ and *X*_*i+1,j*_, are adjacent, their enclosures *ϒ*_*i,j*_ and *ϒ*_*i+1,j*_ are overlapping which ensures a form of continuity in their associated distribution functions. Because *X*_*i,j*_ and *X*_*i+1,j*_ are non-overlapping, each point in the correlation matrix is associated with a unique empirical distribution function. We employ this approach to avoid discontinuities in the normalization functions at the boundary of the submatrices. The size of *X*_*i,j*_ and *ϒ*_*i,j*_ is chosen by the user and controls the degree of smoothing, not unlike bandwidth for a density estimator. In our application, we are using 60 × 60 outer enclosures, with each outer enclosure containing approximately 400 × 400 gene-gene correlations and each inner enclosure containing approximately 200 × 200 gene-gene correlations. We found this setting to work well across our applications.

**Figure 5.**
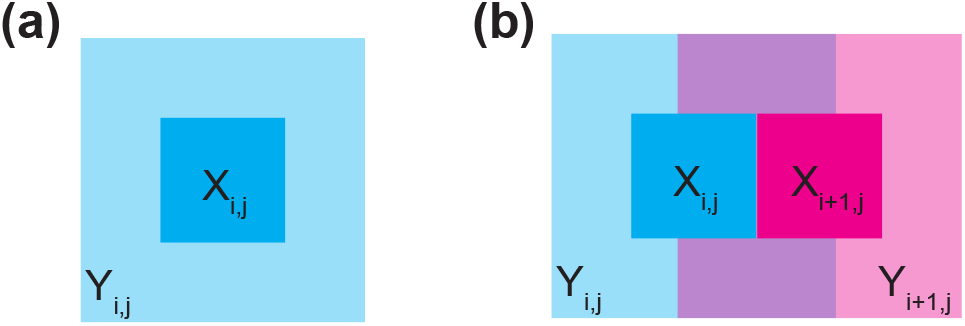
Spatial quantile normalization explained. **(a)** A submatrix *X*_*i,j*_ of the correlation matrix, and its enclosing submatrix *ϒ*_*i,j*_. **(b)** Two directly adjacent, non-overlapping, submatrices *X*_*i,j*_, *X*_*i+1,j*_ and their enclosing, overlapping, submatrices *ϒ*_*i,j*_, *ϒ*_*i+1,j*_. The enclosing submatrices *ϒ*_*i,j*_, *ϒ*_*i+1,j*_ are used to form the empirical distribution functions 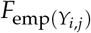 which are then applied to to the non-overlapping submatrices as 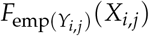.

The choice of reference distribution (with quantile function *q*_target_) can be arbitrary, but in this application we require that its support of the distribution should be contained in [−1, 1]. We recommend a specific submatrix of the correlation matrix to be the reference distribution, specifically the (9, 9)-correlation submatrix (out of a 10×10 binning). This is based on the insights from the preceding section which suggests that the observed pair-wise correlation of two genes is equal to the unobserved pairwise correlation of their expression levels, provided the two genes are highly expressed and thereby less affected by technical noise. We avoid the top right sub-matrix ((10, 10)) because it may contain a wide range of expression levels since we form submatrices of equal size; note the unusual behavior of the top right submatrix in Figure 4. For more details on SpQN, we refer the reader to the Methods section.

Note that mean expression is only used to sort the correlation matrix. We use expression level as a measure of distance between genes so that genes with similar expression level are close (see Discussion for additional comments on the generality of SpQN).

### Spatial quantile normalization removes the mean-correlation relationship

Applying spatial quantile normalization to the 350 RNA-seq samples from adipose subcutaneous tissue in GTEx, we found that it corrects both the background and signal distributions (Figure 6a) across the correlation matrix, thereby removing the mean-correlation relationship (Figure 6b-c). In addition, our normalization method removes the bias towards highly expressed genes when using thresholding to identify highly co-expressed genes (Figure 6d).

**Figure 6.**
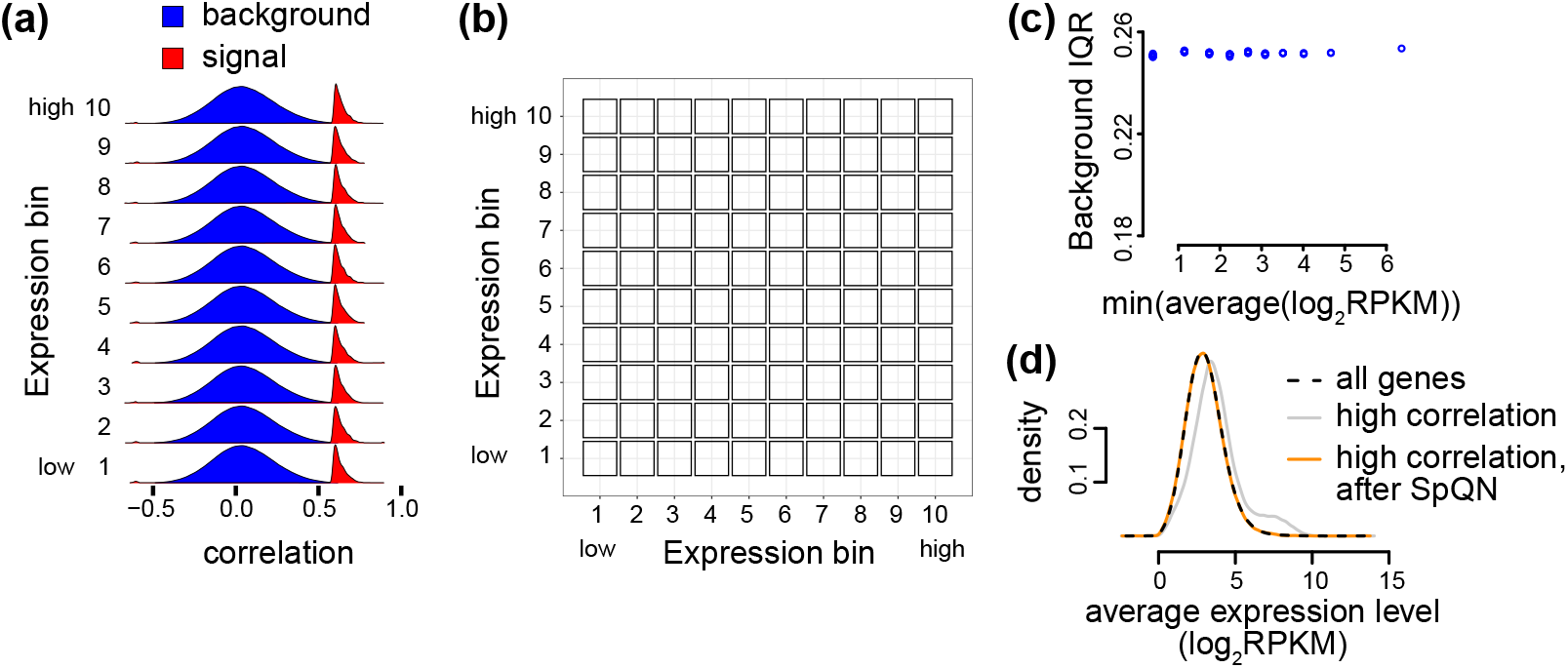
Spatial quantile normalization removes the mean-correlation relationship. Same data as Figure 2, but after applying spatial quantile normalization (SpQN). **(a)** Like Figure 3a, i.e. densities of the Pearson correlation between all genes within each of 10 expression bins (background) as well as the top 0.1% correlations (possible signal). **(b)** Like Figure 2b, i.e. IQRs of Pearson correlations between genes in each of 10 different expression levels. **(c)** Like Figure 2c, i.e. the relationship between IQR of gene-gene correlation distribution and the lowest of the two expression bins associated with the submatrix. **(d)** Like Figure 3b, i.e. the expression level of pairs of genes in different subsets (all genes (black), genes above the 0.1% threshold with (orange) and without SpQN (gray)).

Our observations holds true across a diverse set of GTEx tissues (Figure 7a, Supplemental Figures S1–S2). For each of the 10×10 submatrices of the tissue-specific correlation matrix, we compute an IQR and we consider the distribution of the IQRs before and after applying spatial quantile normalization (Supplemental Figures S1–S2). We observe that using our approach makes the IQRs similar across the correlation submatrices, as the width of the boxplots is smaller for SpQN compared to pre-SpQN (Figure 7a) across all 9 tissues. We note that – because of our choice of target distribution – the IQRs of most of the background distributions increase following spatial quantile normalization. We furthermore note that the pre- and post-SpQN range of IQR is tissue dependent, but this is also driven by differences in sample sizes for the different tissues (Figure 7b).

**Figure 7.**
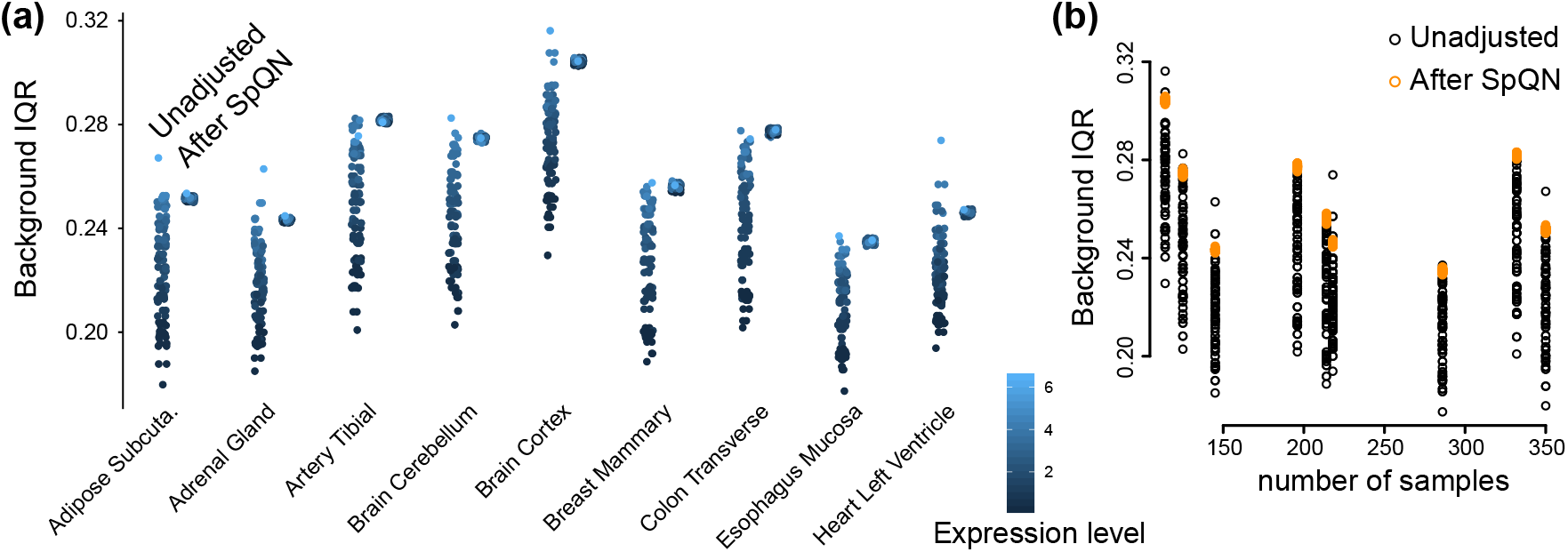
IQR of gene-gene correlation distributions in each bin for 9 tissues. RNA-seq data from GTEx Consortium (2017) from 9 tissues with 4 PCs removed. **(a)** Background IQR for unadjusted (left smear) and SpQN-adjusted (right smear) gene-gene correlation distributions for all expression bins across 9 GTEx tissues. Color indicates expression level. **(b)** The relationship between sample size for a tissue and background IQR for correlation distributions before and after SpQN adjustment.

To highlight the impact of our method on biological relationships, we focus on transcription factors, which have been found to be relatively lowly expressed (Vaquerizas et al., 2009; Boukas et al., 2019). As transcription factors are an important class of regulatory genes, there is substantial interest in identifying co-expression between transcription factors and other genes.

To quantify the impact of our method on transcription factor co-expression, we use a comprehensive list of 1,254 human transcription factors (Barrera et al., 2016). For each of our 9 examplar tissues, we again threshold the correlation matrix and ask how many edges involve transcription factors with and without the use of SpQN. Figure 8 displays the percent increase in edges involving transcription factors following SpQN for various signal thresholds of the correlation matrix. We observe that across all tissues and all signal thresholds, SpQN results in identifying on average 10% more edges involving these genes.

**Figure 8.**
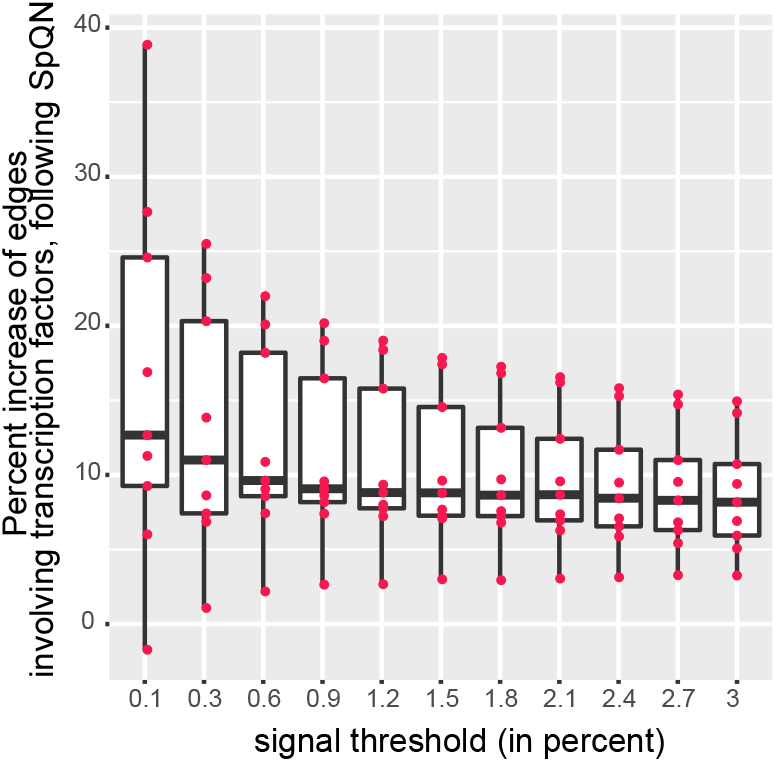
Impact of SpQN on transcription factor co-expression. The precent increase in number of edges (y-axis) identified after thresholding (x-axis) involving transcription factors, following the application of SpQN. Each point is one of 9 tissues from GTEx Consortium (2017). The x-axis shows the thresholding level, i.e. the percentage of gene-gene correlations which are labelled high.

### Mean-correlation bias in single-cell RNA-seq data

Single-cell RNA-sequencing (scRNA-seq) data exhibit the same mean-correlation relationship as in bulk RNA-seq data. We conclude this based on a re-analysis of an unusually deeply sequencing single cell data set Deng et al. (2014). We focus on this particular data set to avoid issues with computing correlation for very sparse data. We kept genes with median log_2_(RPKM) greater than 0, leaving 6,915 out of 22,958 genes.

As depicted in Figure 9, this dataset exhibits the same behavior as the GTEx tissues analyzed above (Figure 9a). Over-representation of highly expressed genes following thresholding the correlation matrix is also observed in this scRNA-seq data (Figure 9b). We observe that the bias of correlation towards highly expressed genes is removed following the application of SpQN (Figure 9b). These observations hold true both when the top 4 PCs are removed (Figure 9a-b) and when we use SVA to estimate the number of PCs (Figure 9c-d), which results in 16 PCs. Unlike the situation for GTEx, we observe that the expression bias is smaller when more PCs are removed (compare Figure 9b to 9d). Together, this suggests that deeply sequenced scRNA-seq data have the same mean-correlation bias as bulk data.

**Figure 9.**
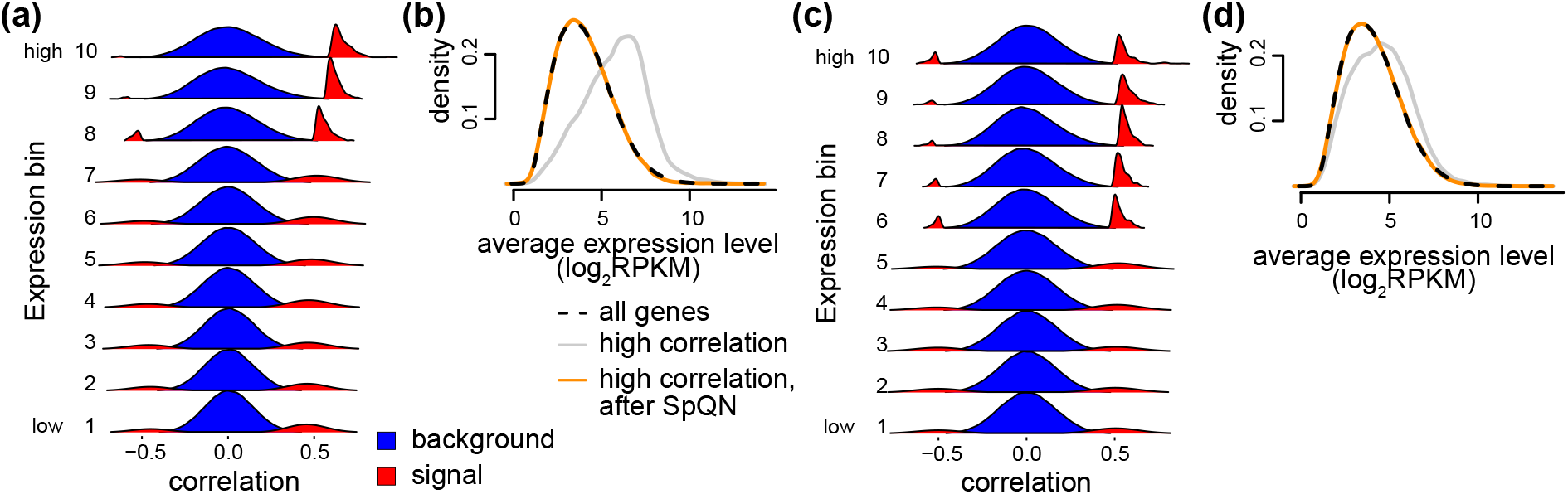
Mean-correlation relationship in scRNA-seq data. scRNA-seq data from Deng et al. (2014) containing 60 cells, with either 4 PCs removed (a-b) or 16 PCs removed (c-d). The later value is the result of using SVA to estimate the number of PCs as suggested by Parsana et al. (2019). **(a)** Like Figure 3a, i.e. densities of the Pearson correlation between all genes within each of 10 expression bins (background, blue) as well as the top 0.1% correlations (possible signal, red). **(b)** Like Figure 6d, i.e. the expression level of pairs of genes in different subsets (all genes (black), genes above the 0.1% threshold with (orange) and without SpQN (gray)). **(c)** Like (a) but for data with 16 PCs removed. **(d)** Like (b) but for data with 16 PCs removed.

### The impact of removing principal components

We now consider the question of the impact of removing unwanted variation in the gene-gene correlation matrix on the mean-correlation relationship.

First, we focus on properties of the data prior to removing top principal components. Considering the background and (possible) signal distributions for all 9 GTEx tissues, we observe that the background distributions are not necessarily centered around zero (Figure 10). Based on our arguments that the background distributions ought to be zero-centered under the assumption of a sparse true correlation network, we term this location shift a “background bias”. The bias is likely batch (but here we depict this as tissues) dependent and so is its relationship with expression level – contrast “Brain Cerebellum” (high bias with high expression) with “Esophagus Mucosa” (high bias with intermediate expression, low bias with both high and low expression). In addition we observe, as previously described, that both the spread of the background distributions as well as the position of the signal distributions are strongly dependent on expression level (and tissue or batch) (Figure 10). Comparing to the same distributions after removing 4 PCs (Figure S1), we conclude that removing even a few PCs has a large beneficial effect on the behaviour of the background distributions, including a substantial reduction in background bias.

**Figure 10.**
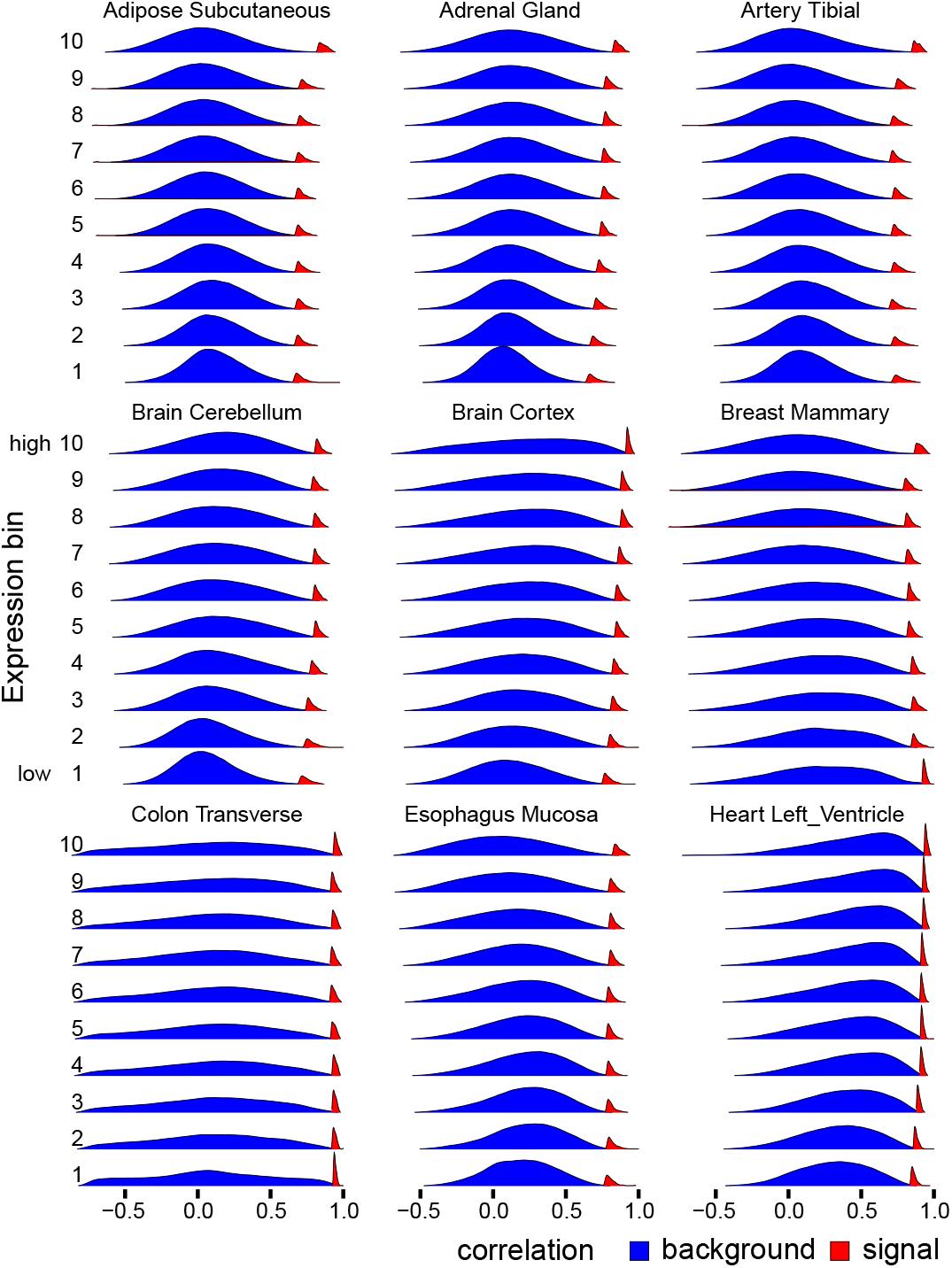
The background and signal components depend on expression level (before removing PCs) Distributions of Pearson correlations for genes within each expression bin, supplemented with the distribution of the top 0.1 % of correlations (within each expression bin).

Next, we focus on removing an increasing number of PCs. If we use the number of PCs recommended by Parsana et al. (2019) (Table 1), we see similar results (Figure 11a) compared to removing 4 PCs (Figure 7a). However, both the average and the spread of the IQRs of background distributions prior to applying spatial quantile normalization, are smaller than using 4 PCs. Interestingly, the variation across tissues in IQRs are now more driven by sample size (Figure 11b) compared to using 4 PCs (Figure 7b).

**Table 1.**
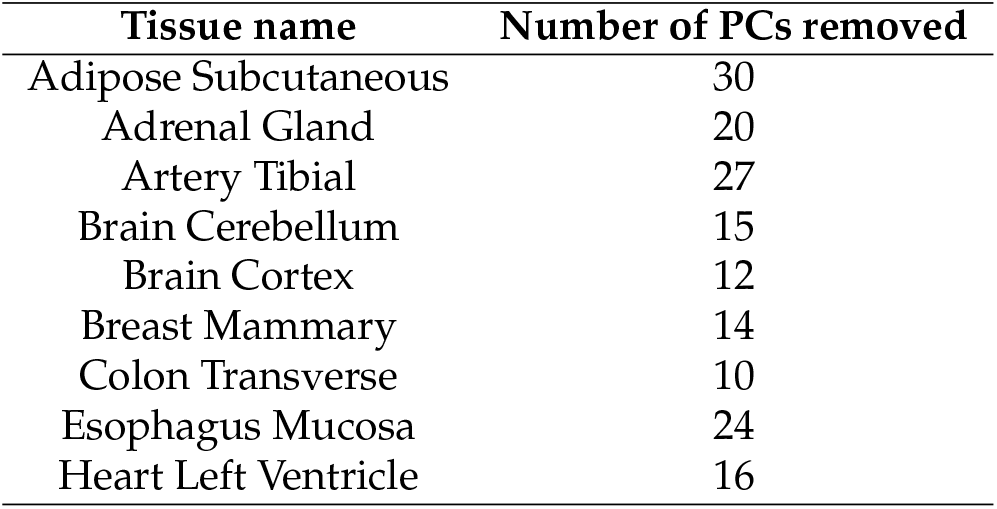
Number of principal components to be removed, as estimated using SVA.

| Tissue name | Number of PCs removed |
| --- | --- |
| Adipose Subcutaneous | 30 |
| Adrenal Gland | 20 |
| Artery Tibial | 27 |
| Brain Cerebellum | 15 |
| Brain Cortex | 12 |
| Breast Mammary | 14 |
| Colon Transverse | 10 |
| Esophagus Mucosa | 24 |
| Heart Left Ventricle | 16 |

**Figure 11.**
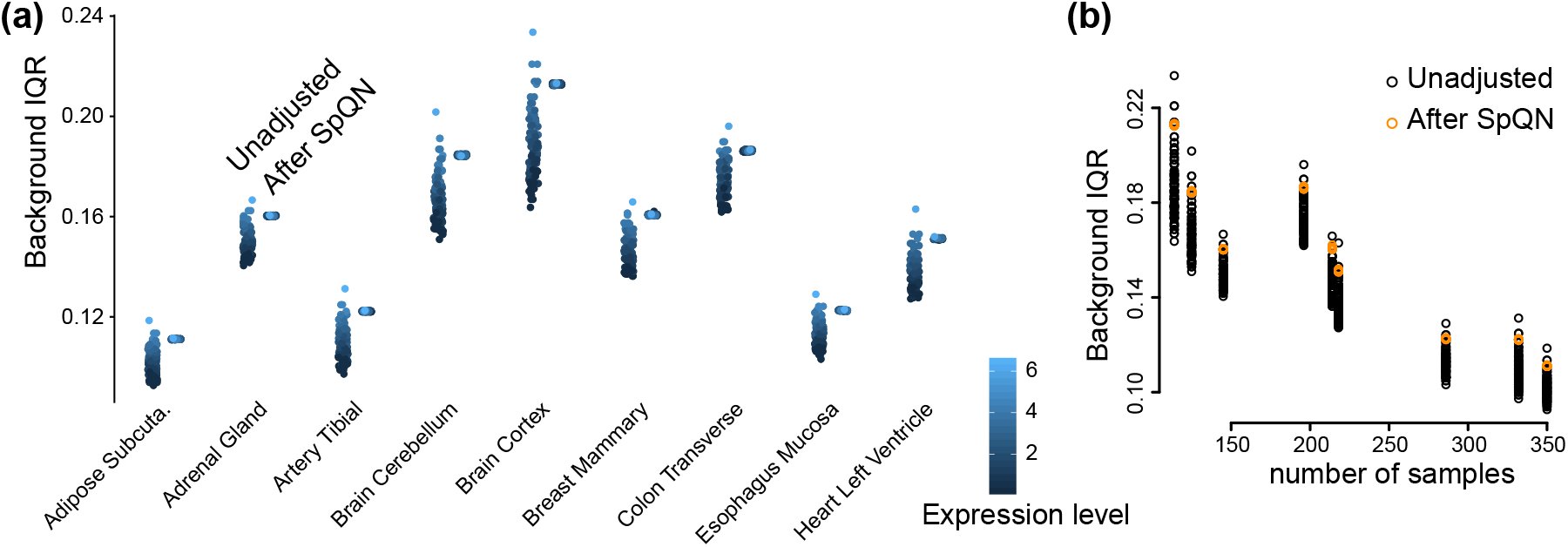
IQR of correlations in each bin for 9 GTEx tissues. Bulk RNA-seq data from GTEx Consortium (2017) from 9 tissues. Each tissue has a number of PCs removed based on the estimate from SVA, as suggested by Parsana et al. (2019). **(a)** Background IQR for unadjusted (left smear) and SpQN-adjusted (right smear) gene-gene correlation distributions for all expression bins across 9 GTEx tissues. Color indicates expression level. **(b)** The relationship between sample size and IQR for correlations before and after adjustment.

Finally, we quantified the effect of removing PCs on the bias and spread of the background distributions (Figure 12). Bias is reduced across all tissues, although the largest decrease happens with the first few PCs (perhaps up to 10 PCs). The spread of the background distributions are also reduced, although for some tissues there is an inflection point after which the spread increases with higher number of PCs. Together, these observations suggests that removing a high number of PCs may have a substantial positive impact on the mean-correlation relationship.

**Figure 12.**
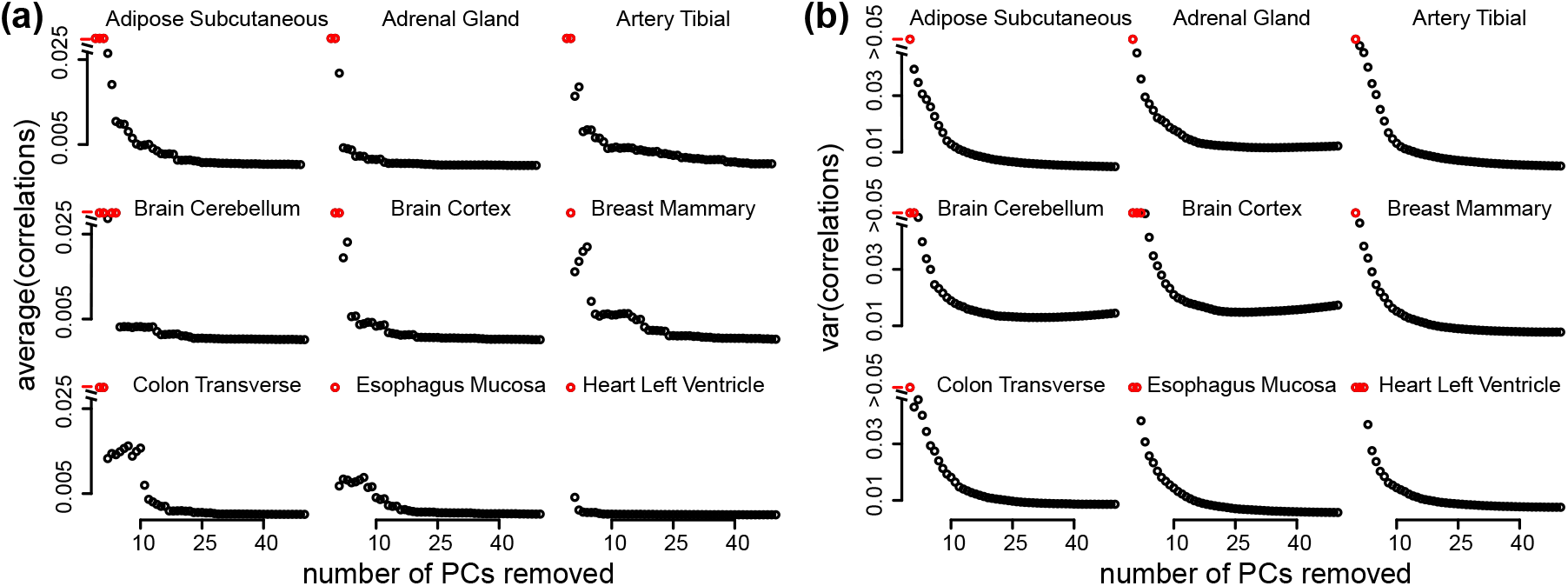
The effect of removing principal components on bias and variance of the background distribution.

**(a)** Average bias, defined as the average of the median of the 10 background distributions. **(b)** Average variance, defined as the average variance of the 10 background distributions. Red colored points have bias or variance exceeding the limits of the plot.

However, more important is the impact on down-stream analysis, particularly the identification of highly co-expressed genes in co-expression analysis. Using our previously described approach of thresholding the correlation matrix, we observe that removing a higher number of principal components has a small impact on the expression bias for highly co-expressed genes (Figure 13), suggesting that focusing on background spread by itself is irrelevant, but that it is more important to evaluate impact on the background distributions relative to the impact on the (possible) signal distributions. Importantly, we observe that spatial quantile normalization removes this expression bias, irrespective of how many principal components were removed.

**Figure 13.**
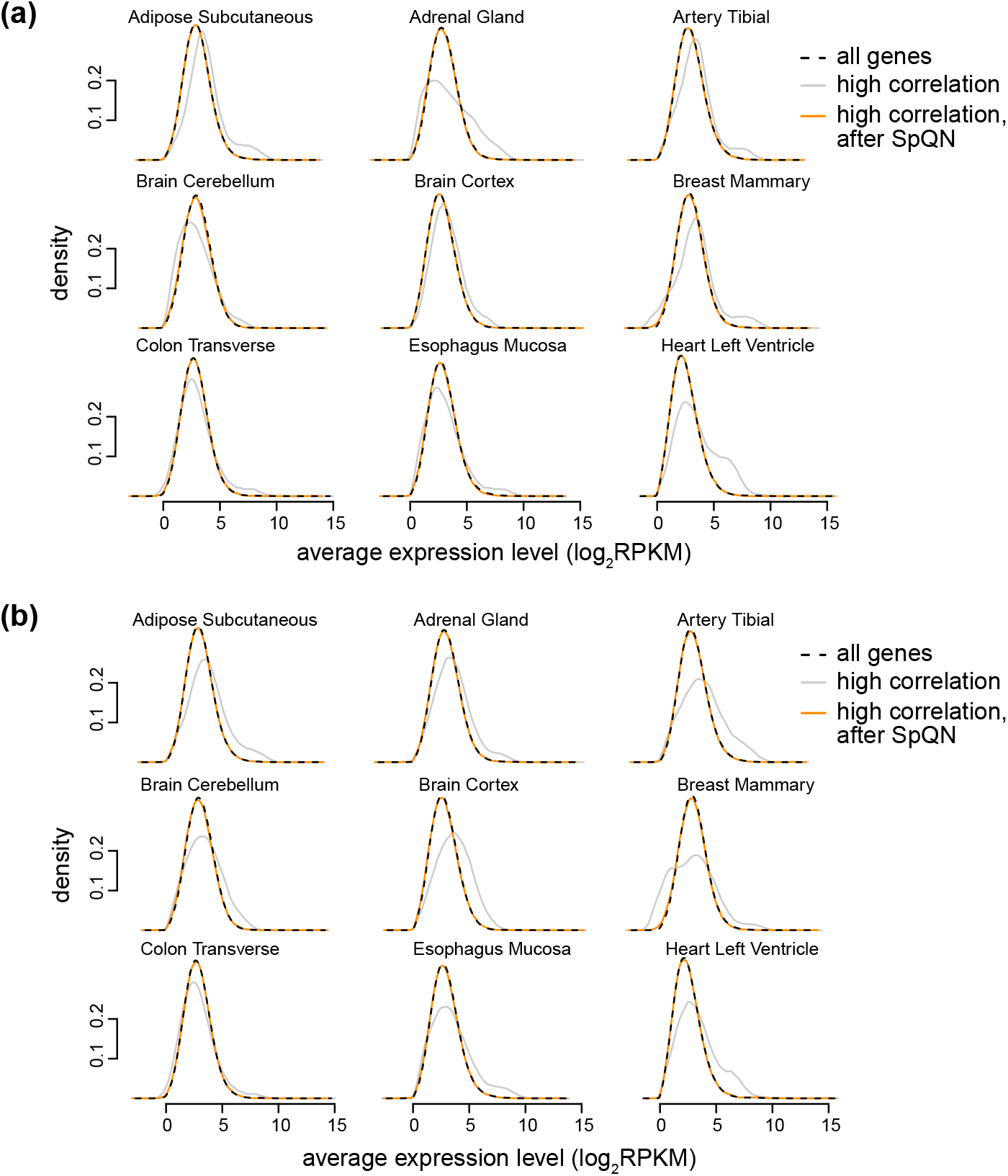
The effect of removing principal components (PCs) on bias towards highly expressed genes. As Figure 6d but for 9 tissues and two different approaches for removing principal components. **(a)** 4 PCs were removed from the correlation matrix. **(b)** SVA was used to estimate the number of PCs to remove (range: 10-30 PCs).

## Discussion

A key challenge in gene network reconstruction methods is to select biologically meaningful pairs of co-expressed genes. A standard approach is to select for highly correlated gene pairs (possibly signal) compared to lowly correlated gene pairs (background). Here, we demonstrate the existence of a *mean-correlation relationship*, which can bias co-expression analysis resulting in an over-representation of highly expressed genes amongst connected gene pairs. This is particularly problematic for genes that are generally expressed at low levels, such as transcription factors. To address this, we have developed a normalization method for the gene-gene correlation matrix that can standardize “local” distributions of the correlations across the matrix. Using nine GTEx tissues with bulk RNA-seq, as well as one deeply sequenced scRNA-seq dataset, we have illustrated how the mean-correlation relationship can be removed from the correlation matrix using spatial quantile normalization (SpQN). Utilizing our method results in a greater number of connections involving transcription factors, an important class of regulatory genes.

Here we have used SpQN for the specific problem of removing the confounding effect of mean expression on the correlation matrix. We highlight two important points about this method. First, the formulation (and software) merely requires the correlation matrix to be sorted according to the confounding variable; the fact that the confounding variable is mean expression is not critical. Second, the core ideas of SpQN extends far beyond correlation matrices. Specifically, the way we propose to use neighbourhoods of different size to define inner and outer enclosures, only requires some sense of distance to define the neighbourhoods. This implies the idea of SpQN can be applied to any kind of data with a natural sense of distance between observations, such as time series data or spatial data.

We have investigated the impact of removing principal components on the mean-correlation relationship, which has revealed a number of interesting observations. First, importantly, while removing principal components impacts the spread and bias of the background distributions, we observe a similar bias towards highly expressed genes when removing 4 principal components as when removing 10-30 principal components. Second, the different patterns of expression dependence in the background distributions is likely to represent tissue-dependent batch effects. This therefore serves to illustrate how batch effects impact co-expression analysis. An interesting question is what forces create these different patterns. Altogether, our work reinforces the message from Parsana et al. (2019) that removing principal components is good practice, although exactly how many components to remove is still an open question.

Together, this work shows the importance of assessing and addressing mean-correlation bias in co-expression analysis, and provides the first method for doing so.

## Methods

### GTEx bulk RNA-seq dataset

We used GTEx dataset for 9 tissues (GTEx Consortium, 2017), including adipose subcutaneous, adrenal gland, artery tibial, brain cerebellum, brain cerebellum, brain cortex, breast mammary, colon transverse, esophagus mucosa and heart left ventricle. The read counts data were downloaded from GTEx portal, release v6p, expression measures `GTEx_Analysis_v6p_RNA-seq_RNA-SeQCv1.1.8_gene_reads.gct.gz`, gene annotation file `gencode.v19.genes.v6p_model.patched_contigs.gtf.gz` and sample information file `GTEx_Data_V6_Annotations_SampleAttributesDS.txt`. We kept protein coding genes and long non-coding genes, and transformed the read counts into log-RPKMs scale using

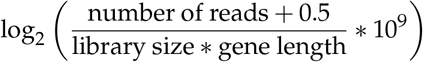

where library size is the total number of reads in the sample. For each tissue, we only kept the genes with median log_2_(RPKM) above 0.

### Single-cell RNA-seq dataset

We downloaded the scRNA-seq data from Deng et al. (2014). RPKM normalized expression matrix of 60 mid-blast cells are used. We transformed the expression matrix with log_2_(RPKM + 0.5), and kept the genes with median log_2_(RPKM + 0.5) greater than 0.

### PPI database

We downloaded PPI data from HuRI database, which contains around 53,000 pairs of protein-protein interactions in human (Luck et al., 2019), with each protein annotated by the corresponding gene ensemble IDs. For each tissue, we only kept the genes overlapped with the genes in the filtered counts.

### Remove batch effect and generate gene correlation matrix

We applied principal component analysis to address the confounding noise from batch effects to the expression matrix through removing leading PCs (Parsana et al., 2019). For each tissue, we scaled the expression matrix such that the expression of each gene has mean 0 and variance 1 across the samples. We regressed out the top PCs and created a matrix of the regression residuals using the WGCNA R package (Langfelder, Horvath, 2008).

For each tissue, we considered two approaches to remove PCs from the correlation matrix. In the first approach, we regressed out the top 4 PCs. In the second approach, we regressed out the number of top PCs determined from using the `num.sv` function in the sva package (Leek, Johnson, et al., 2012). Based on the residuals from these matrices, we generated the gene-gene correlation matrix by calculating the Pearson correlation coefficient of the residuals between genes, for each type of residuals and every tissue.

### A model for correlation in RNA-seq data

Let *ϒ*_1_, *ϒ*_2_ be the data from two observed genes and *Z*_1_, *Z*_2_ be the unobserved “true” expression variables. We assume that E(*ϒ* | *Z*) = *Z*. If we furthermore assume that *ϒ*_1_ and *ϒ*_2_ are conditional independent given (*Z*_1_, *Z*_2_) (independence of the sequencing noise) we get

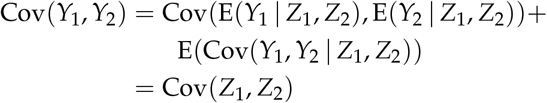

because of our two assumptions. By multiplying and dividing with the same factors we now immediately get

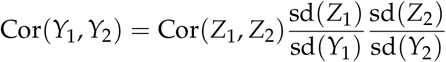

Looking at the standard deviations of the expression level, we get

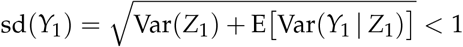

If we make the additional assumption of Var(*ϒ*_1_ | *Z*_1_) = *Z*_1_, we obtain

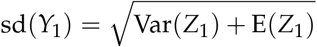

Forming the ratio and dividing both the numerator and the denominator by 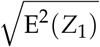 yields

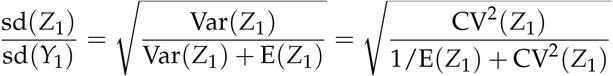

These assumptions hold when *ϒ*_1_ |*Z*_1_ ~ Poisson(*Z*_1_) and can easily be modified to include non-random library sizes. However, we apply these results to log_2_(RPKM) (including after removal of principal components), and it is worthwhile to consider these assumptions in general. The first assumption, that E(*ϒ | Z*) = *Z* is necessary to recover the correlation, but can easily be relaxed to the conditional expectation being an affine function of *Z*; E(*ϒ | Z*) = *α* + *βZ* where *α* will disappear and *β* be part of the adjustment factor. The second assumption, Var (*ϒ*_1_ | *Z*_1_) = *Z*_1_, is unnecessary – the main requirement is that E[Var(*ϒ*_1_ | *Z*_1_)/E^2^(*Z*_1_) vanishes when E(*Z*_1_) is large.

### Spatial quantile normalization

To correct the mean-correlation bias in the gene correlation matrix, we developed a method called spatial quantile normalization (SpQN) that removes the difference in “local” distribution of correlations across the gene correlation matrix.

Binning the correlation matrix into disjoint grids and normalizing them separately could result in artifacts. The within-bin variance of distribution could result in imprecise estimation of the local distribution, and therefore the normalization would lead to high within-bin variance. On the other hand, using small bins could result in insufficient sample size for approximating local distribution. We therefore introduce a quantile normalization method that could address this binning problem.

For each tissue, we used the expression matrix (on the log_2_ (RPKM) scale), with the genes sorted according to the expression level (average expression across samples). We grouped the genes into 10 separate disjoint bins and numbered them by 1,2,…,10, with the expression level from low to high. For the corresponding correlation matrix with the genes sorted in the same way, we stratified the matrix into 10 by 10 disjoint equal-size grids, and numbered the grids as (*i*, *j*) using the expression level for the *i*^*th*^ and *j*^*th*^ bin. The selection of reference distribution, *F*_ref_, can be arbitrary, but we used the empirical distribution of grid (9,9).

The set of disjoint submatrices *X*_*i,j*_ and the larger submatrices *ϒ*_*i,j*_ that embeds *X*_*i,j*_ were assigned based on the preset parameters – number of bins (written as *n*_*group*_) and size of larger bins (written as *w*), with default settings *n*_*group*_=60 and *w*=400.

{*ϒ*_*i,j*_} is assigned to be equal-size, equal-distance and overlapped bins that covered the correlation matrix, with bin size *w* and distance *d* = (*n*_gene_ – *w*)/(*n*_*group*_ – 1), where *n*_gene_ is the number of genes. {*ϒ*_*i,j*_} can be written as

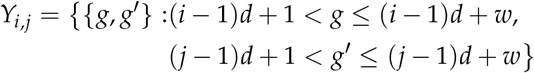

for *i*, *j* = 1, 2, …, *n*_group_.

The set of submatrices {*X*_*i,j*_} is assigned to be disjoint and same-distance bins, with distance equals to that of *ϒ*_*i,j*_, written as

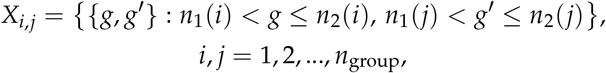

where

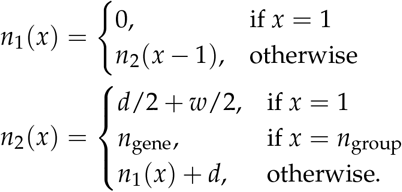

Using using the empirical distribution of *ϒ*_*i,j*_, we estimated local distribution for *X*_*i,j*_,

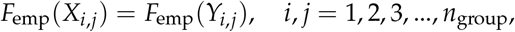

We mapped the correlations within each disjoint bin to the corresponding quantiles in the reference distribution,

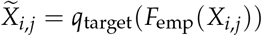

where *q*_target_ is the quantile in the reference distribution, and 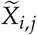 is the correlations in *X*_*i,j*_ after quantile normalization. If *ϒ*_*i,j*_ is the same as *X*_*i,j*_ this is the same as quantile normalization. As *ϒ*_*i,j*_ gets larger than *X*_*i,j*_, *F*_emp_(*X*_*i,j*_) is only approximately uniform and the corrected correlations 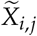 are only approximately following the reference distribution.

## Software availability

Analysis scripts and an R package for spatial quantile normalization is available at https://github.com/hansenlab/spqn.

## Supplementary Materials

**Supplementary Figure S1.**
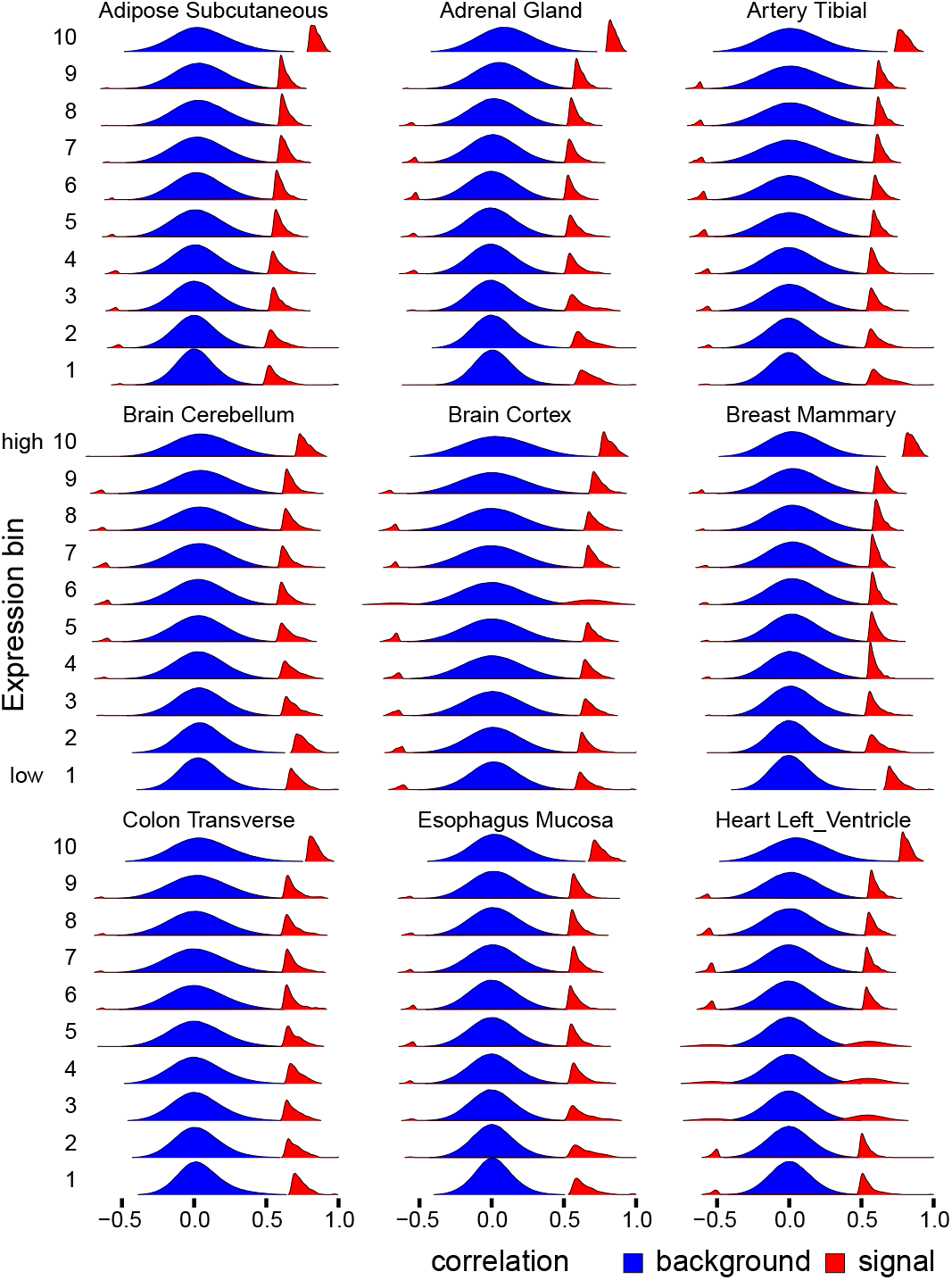
The background and signal components depends on expression level across many tissues. Data is 9 different GTEx tissues, all with 4 PCs removed. Distributions of Pearson correlations for genes within each expression bin, supplemented with the distribution of the top 0.1 % of correlations (within each expression bin).

**Supplementary Figure S2.**
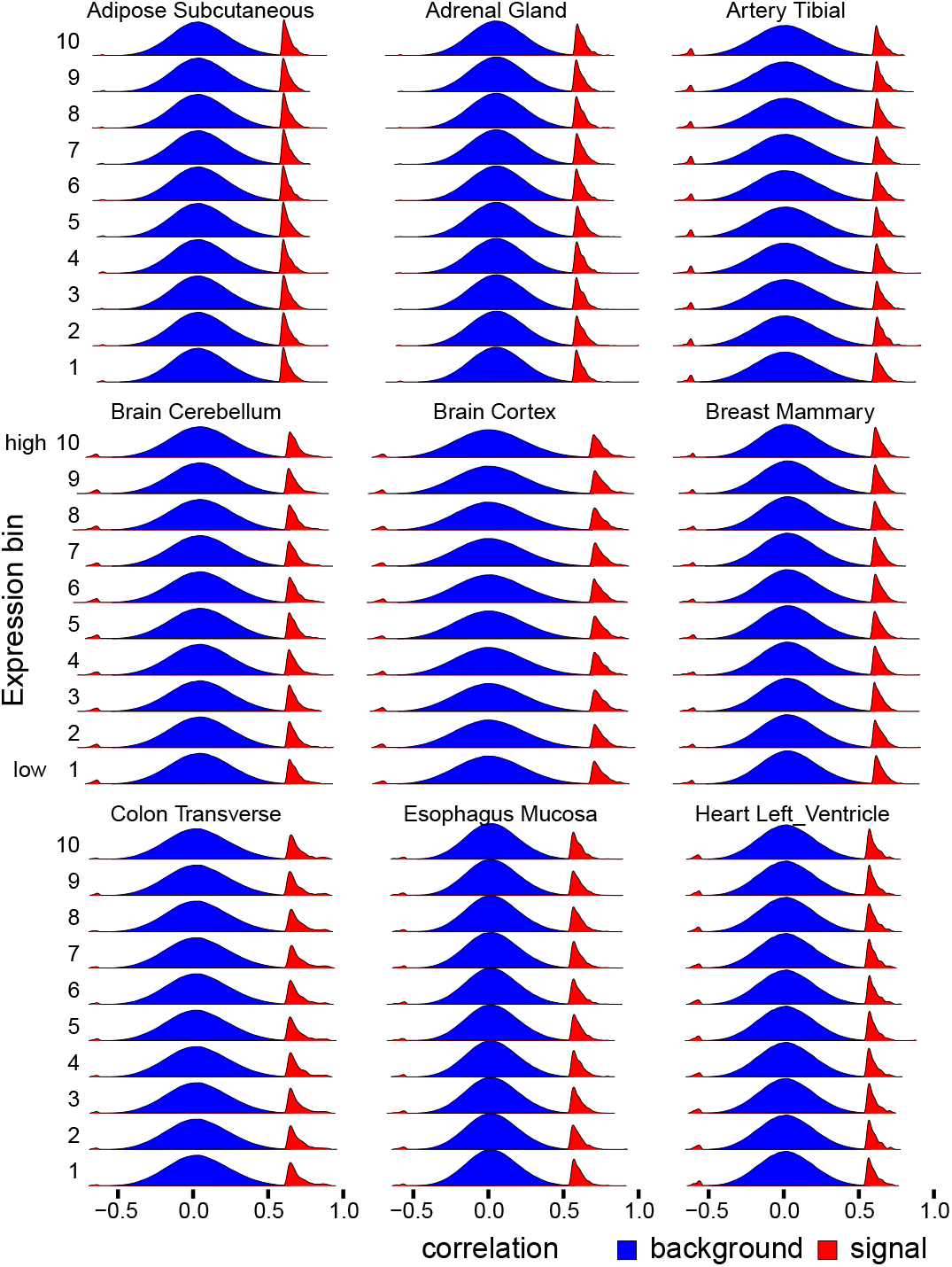
The background and signal components depends on expression level after spatial quantile normalization. Data is 9 different GTEx tissues, all with 4 PCs removed. Like Supplemental Figure S1 but after applying spatial quantile normalization.

## References

Amaratunga D, Cabrera J. 2001. Analysis of Data from Viral DNA Microchips. Journal of American Statistical Association 96.456: 1161. D O I: 10.1198/016214501753381814.

Barrera LA, Vedenko A, Kurland JV, Rogers JM, Gisselbrecht SS, Rossin EJ, Woodard J, Mariani L, Kock KH, Inukai S, et al. 2016. Survey of variation in human transcription factors reveals prevalent DNA binding changes. Science 351.6280: 1450–1454. D O I: 10.1126/science.aad2257.

Bolstad B, Irizarry R, Astrand M, Speed T. 2003. A comparison of normalization methods for high density oligonucleotide array data based on variance and bias. Bioinformatics 19.2: 185–193. D O I: 10.1093/bioinformatics/19.2.185.

Boukas L, Havrilla JM, Hickey PF, Quinlan AR, Bjornsson HT, Hansen KD. 2019. Coexpression patterns define epigenetic regulators associated with neurological dysfunction. Genome Research 29.4: 532–542. D O I: 10.1101/gr.239442.118.

Bullard JH, Purdom E, Hansen KD, Dudoit S. 2010. Evaluation of statistical methods for normalization and differential expression in mRNA-Seq experiments. BMC Bioinformatics 11: 94. D O I: 10.1186/1471-2105-11-94.

Deng Q, Ramsköld D, Reinius B, Sandberg R. 2014. Single-cell RNA-seq reveals dynamic, random monoallelic gene expression in mammalian cells. Science 343.6167: 193–196. D O I: 10.1126/science.1245316.

Farahbod M, Pavlidis P. 2019. Differential coexpression in human tissues and the confounding effect of mean expression levels. Bioinformatics 35.1: 55–61. D O I: 10.1093/bioinformatics/bty538.

Freytag S, Gagnon-Bartsch J, Speed TP, Bahlo M. 2015. Systematic noise degrades gene co-expression signals but can be corrected. BMC Bioinformatics 16: 309. D O I: 10.1186/s12859-015-0745-3.

Friedman J, Hastie T, Tibshirani R. 2008. Sparse inverse co-variance estimation with the graphical lasso. Biostatistics 9.3: 432–441. D O I: 10.1093/biostatistics/kxm045.

Ghazalpour A, Doss S, Zhang B, Wang S, Plaisier C, Castellanos R, Brozell A, Schadt EE, Drake TA, Lusis AJ, et al. 2006. Integrating genetic and network analysis to characterize genes related to mouse weight. PLOS Genetics 2.8: e130. D O I: 10.1371/journal.pgen.0020130.

GTEx Consortium. 2017. Genetic effects on gene expression across human tissues. Nature 550.7675: 204–213. D O I: 10.1038/nature24277.

Langfelder P, Horvath S. 2008. WGCNA: an R package for weighted correlation network analysis. BMC Bioinformatics 9: 559. D O I: 10.1186/1471-2105-9-559.

Leek JT, Johnson WE, Parker HS, Jaffe AE, Storey JD. 2012. The sva package for removing batch effects and other unwanted variation in high-throughput experiments. Bioinformatics 28.6: 882–883. D O I: 10.1093/bioinformatics/bts034.

Leek JT, Storey JD. 2007. Capturing heterogeneity in gene expression studies by surrogate variable analysis. PLOS Genetics 3.9: 1724–1735. D O I: 10.1371/journal.pgen.0030161.

Leek JT, Storey JD. 2008. A general framework for multiple testing dependence. Proceedings of the National Academy of Sciences of the United States of America 105.48: 18718–18723. D O I: 10.1073/pnas.0808709105.

Luck K, Kim DK, Lambourne L, Spirohn K, Begg BE, Bian W, Brignall R, Cafarelli T, Campos-Laborie FJ, Charloteaux B, et al. 2019. A reference map of the human protein interactome. bioRxiv: 605451. D O I: 10.1101/605451.

Marioni JC, Mason CE, Mane SM, Stephens M, Gilad Y. 2008. RNA-seq: an assessment of technical reproducibility and comparison with gene expression arrays. Genome Research 18.9: 1509–1517. D O I: 10.1101/gr.079558.108.

Oldham MC, Konopka G, Iwamoto K, Langfelder P, Kato T, Horvath S, Geschwind DH. 2008. Functional organization of the transcriptome in human brain. Nature Neuroscience 11.11: 1271–1282. D O I: 10.1038/nn.2207.

Parsana P, Ruberman C, Jaffe AE, Schatz MC, Battle A, Leek JT. 2019. Addressing confounding artifacts in reconstruction of gene co-expression networks. Genome Biology 20.1: 94. D O I: 10.1186/s13059-019-1700-9.

Saha A, Kim Y, Gewirtz ADH, Jo B, Gao C, McDowell IC, GTEx Consortium, Engelhardt BE, Battle A. 2017. Co-expression networks reveal the tissue-specific regulation of transcription and splicing. Genome Research 27.11: 1843–1858. D O I: 10.1101/gr.216721.116.

Vaquerizas JM, Kummerfeld SK, Teichmann SA, Luscombe NM. 2009. A census of human transcription factors: function, expression and evolution. Nature Reviews Genetics 10.4: 252–263. D O I: 10.1038/nrg2538.

Willsey AJ, Sanders SJ, Li M, Dong S, Tebbenkamp AT, Muhle RA, Reilly SK, Lin L, Fertuzinhos S, Miller JA, et al. 2013. Coexpression networks implicate human midfetal deep cortical projection neurons in the pathogenesis of autism. Cell 155.5: 997–1007. D O I: 10.1016/j.cell.2013.10.020.

Workman C, Jensen LJ, Jarmer H, Berka R, Gautier L, Nielser HB, Saxild HH, Nielsen C, Brunak S, Knudsen S. 2002. A new non-linear normalization method for reducing variability in DNA microarray experiments. Genome Biology 3.9: research0048. DOI: 10.1186/gb-2002-3-9-research0048.

Zhang Y, Cuerdo J, Halushka MK, McCall MN. 2019. The effect of tissue composition on gene co-expression. Briefings in Bioinformatics: bbz135. DOI: 10.1093/bib/bbz135.

